# Inhibitory co-transmission from midbrain dopamine neurons relies on presynaptic GABA uptake

**DOI:** 10.1101/2021.11.26.470142

**Authors:** Riccardo Melani, Nicolas X. Tritsch

## Abstract

Dopamine (DA)-releasing neurons in the substantia nigra pars compacta (SNc^DA^) inhibit target cells in the striatum through postsynaptic activation of γ-aminobutyric acid (GABA) receptors. However, the molecular mechanisms responsible for GABAergic signaling remain unclear, as SNc^DA^ neurons lack enzymes typically required to produce GABA or package it into synaptic vesicles. Here we show that aldehyde dehydrogenase 1a1 (Aldh1a1), an enzyme proposed to function as a GABA synthetic enzyme in SNc^DA^ neurons does not produce GABA for synaptic transmission. Instead, we demonstrate that SNc^DA^ axons obtain GABA exclusively through presynaptic uptake using the membrane GABA transporter Gat1 (encoded by *Slc6a1*). GABA is then packaged for vesicular release using the vesicular monoamine transporter Vmat2. Our data therefore show that presynaptic transmitter recycling can substitute for *de novo* GABA synthesis and that Vmat2 contributes to vesicular GABA transport, expanding the range of molecular mechanisms available to neurons to support inhibitory synaptic communication.

## INTRODUCTION

A detailed appreciation for how neurons modify the activity of surrounding cells through the release of small-molecule and peptide transmitters is key to our understanding of the brain. Because neurons rely primarily on the release of a single classical neurotransmitter like acetylcholine, glutamate or γ-aminobutyric acid (GABA) to transmit electrical signals across synapses, it is typically sufficient to manipulate the activity of presynaptic cells to evaluate the postsynaptic contribution of individual transmitters. However, this approach is insufficient for many neurons within the central nervous system that release multiple classical transmitters. Chief amongst them are ventral midbrain neurons of the substantia nigra pars compacta (SNc), which are critical for the execution and learning of goal-directed movements, and notorious for their propensity to degenerate in Parkinson’s disease. These cells are best known for their ability to release the neuromodulator dopamine (DA), but they can also directly affect the discharge of target cells in dorsal striatum through mono-synaptic glutamatergic and GABAergic signaling (Hnasko et al., 2010; Tecuapetla et al., 2010; Tritsch et al., 2012; Trudeau et al., 2014). The molecular mechanisms that enable glutamate co-release from DA-releasing SNc (SNc^DA^) neurons are well established (Zhang et al., 2015; Eskenazi et al., 2021; Hnasko and Edwards, 2011). By contrast, little is known about inhibitory co-transmission, despite the fact that SNc^DA^ neurons strongly inhibit striatal output through large monosynaptic GABA_A_ receptor-mediated currents in spiny projection neurons (SPNs) (Tritsch et al., 2012, 2014; Nelson et al., 2014; Kim et al., 2015; Dorst et al., 2020; Ishikawa et al., 2013).

An important contributor to this gap in knowledge is that nigrostriatal DAergic neurons do not possess the molecular machinery classically thought to be essential for GABAergic synaptic transmission; they do not express the GABA synthetic enzymes Gad65 and Gad67 (encoded by *Gad2* and *Gad1*, respectively), or the vesicular GABA transporter Vgat (also known as Viaat or Slc32a1) (Tritsch et al., 2012, 2014; Poulin et al., 2014; Kim et al., 2015; Saunders et al., 2018). Instead, inhibitory co-transmission from SNc^DA^ neurons relies on the vesicular monoamine transporter Vmat2 (encoded by *Slc18a2*) (Tritsch et al., 2012), which is not known to transport GABA (Yelin and Schuldiner, 1995). Importantly, several endogenous ligands activate GABA_A_ receptors in addition to GABA, including taurine, β-alanine and the histamine metabolite imidazole-4-acetic acid (Johnston, 1996), and GABA_A_ receptor-mediated currents are strongly modulated by neurosteroids like allopregnanolone (Belelli and Lambert, 2005). It is therefore conceivable that the inhibitory neurotransmitter released by SNc^DA^ neurons is not GABA itself, but a novel molecule that functions as a GABA_A_ receptor agonist or allosteric modulator and that serves as a substrate for Vmat2, such as naturally-occurring analogs of the synthetic GABA_A_ receptor agonists muscimol, THIP (4,5,6,7-tetrahydroisoxazolo[5,4-c]pyridin-3-ol) and isoguvacine (Johnston, 1996). Another possibility is that GABA is the transmitter that SNc^DA^ neurons release, but that the mechanisms involved remain to be defined. Both scenarios point to an incomplete understanding of the molecular pathways that neurons can leverage to inhibit postsynaptic targets.

We previously suggested that SNc^DA^ neurons import GABA from the extracellular milieu, as they contain mRNA for the plasma membrane GABA transporter Gat1 (encoded by *Slc6a1*) and pharmacological inhibition of extracellular GABA uptake in striatum impairs GABAergic co-transmission (Tritsch et al., 2014). However, a recent study found that Gat1 does not distribute to the axon of SNc^DA^ neurons and that Gat1 inhibition non-selectively depresses vesicular release from SNc^DA^ axons by elevating extracellular GABA levels in striatum (Roberts et al., 2020). In addition, SNc^DA^ neurons were proposed to synthetize GABA *de novo* using a non-conventional mitochondrial pathway dependent on the enzyme aldehyde dehydrogenase 1a1 (Aldh1a1) (Kim et al., 2015), which a subpopulation of SNc^DA^ neurons express abundantly (McCaffery and Dräger, 1994; Galter et al., 2003; Liu et al., 2014; Poulin et al., 2014, 2018). Other neural and non-neural cells expressing this and related aldehyde dehydrogenases have consequently been proposed to employ a similar mechanism to produce GABA and influence brain function (Carmichael et al., 2021; Kwak et al., 2020; Jin et al., 2021).

Our study identifies the mechanism that enables inhibitory neurotransmission from SNc^DA^ neurons. Using a combination of genetic and molecular manipulations, optogenetics and whole-cell electrophysiological recordings in mice, we demonstrate that GABA is the neurotransmitter that SNc^DA^ neurons co-release in striatum. We find that Aldh1a1 does not produce GABA, and show that SNc^DA^ neurons rely instead on Gat1-mediated transmembrane uptake to obtain GABA for Vmat2-dependent vesicular release. Our data therefore show that membrane GABA transporters and vesicular monoamine transporters can respectively substitute for Gad-dependent GABA synthesis and Vgat-mediated vesicular loading, expanding the range of molecular mechanisms available to neurons to support inhibitory neurotransmission.

## RESULTS

### Aldh1a1 affects inhibitory co-transmission from SNc^DA^ neurons in a non-cell autonomous fashion

Aldh1a1 is a multifunctional protein highly expressed in ventral tier SNc^DA^ neurons that innervate parts of the dorsal striatum (Figures 1A and 1B) (Carmichael et al., 2021; McCaffery and Dräger, 1994; Poulin et al., 2014, 2018; Wu et al., 2019). Recent evidence suggests that Aldh1a1 acts as a mitochondrial GABA synthetic enzyme that provides the majority of the GABA that SNc^DA^ neurons release onto SPNs (Kim et al., 2015). To examine this, we obtained Aldh1a1 germline knockout mice (*Aldh1a1*^-/-^; Fan et al., 2003). Using immunofluorescence, we confirmed that Aldh1a1 protein is no longer detected in the brain of knockouts, including in ventral tier SNc^DA^ neurons and their axons in striatum (Figures 1A and 1B).

**Figure 1.**
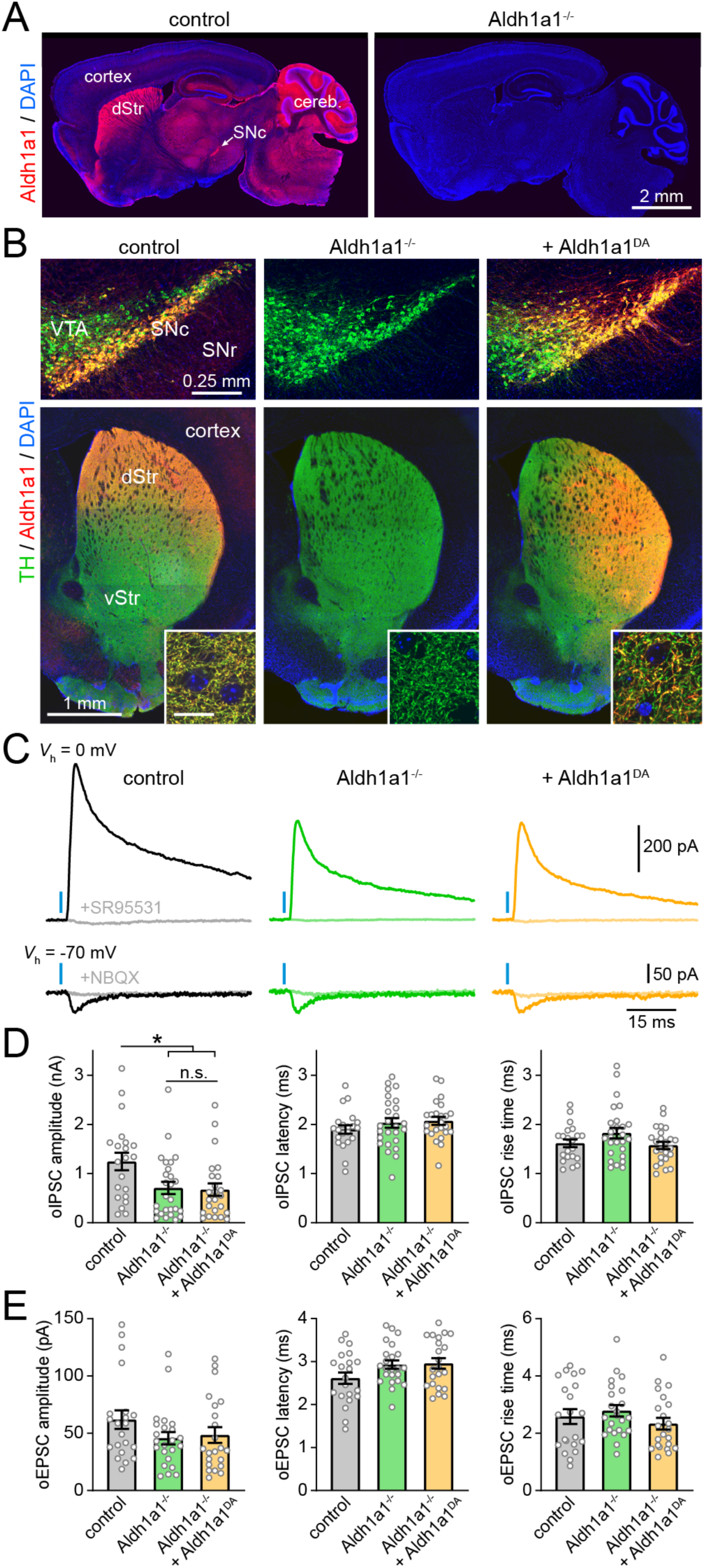
Non-cell autonomous modulation of GABA co-release by Aldh1a1. **A)** Sagittal brain sections from control (left) and Aldh1a1^-/-^ (right) mice immunolabeled for Aldh1a1 (red) and DAPI nuclear stain (blue). Aldh1a1 is widely expressed throughout the central nervous system, including in dorsal striatum (dStr)-projecting SNc^DA^ neurons. Cereb: cerebellum. **B)** Coronal brain sections from a control *Dat*^IRES-Cre/+^ mouse (left), a *Dat*^IRES-Cre/+^;Aldh1a1^-/-^ mouse (middle) and a *Dat*^IRES-Cre/+^;Aldh1a1^-/-^ mouse in which midbrain DA neurons are selectively transduced with Cre-dependent Aldh1a1 (right) immunolabeled for tyrosine hydroxylase (TH, green), Aldh1a1 (red) and DAPI nuclear stain (blue). Top row: confocal image of ventral midbrain showing SNc, ventral tegmental area (VTA) and substantia nigra pars reticulata (SNr). Bottom row: Epifluorescence image of striatum. vStr: ventral striatum. Inset: confocal image of dorsal striatum confirming Aldh1a1 distribution in axons (scale bar: 10 µm). **C)** Example postsynaptic current responses recorded from SPNs voltage-clamped (*V*_h_) at the reversal potential for glutamatergic currents (top) or GABAergic currents (bottom) upon optogenetic stimulation of SNc^DA^ axons (blue bar) before and after bath application of the GABA_A_ receptor antagonist SR95531 (10 µM) or the ionotropic glutamate receptor blockers NBQX (10 µM) in the same experimental conditions as in (**B**). **D)** Quantification of optogenetically-evoked IPSC amplitude (left), synaptic latency (middle) and 10-90% rise time (right) in slices from *Dat*^IRES-Cre/+^ (control), *Dat*^IRES-Cre/+^;Aldh1a1^-/-^ and *Dat*^IRES-Cre/+^;Aldh1a1^-/-^ mice expressing Aldh1a1 only in SNc^DA^ neurons. Kruskall-Wallis tests revealed a significant effect for oIPSC amplitude (p = 0.014), but not latency (p=0.38) or rise time (p = 0.23). * p = 0.025 between control and Aldh1a1^-/-^, and p = 0.018 between control and Aldh1a1^-/-^ + Aldh1a1^DA^, Dunn’s Multiple Comparison. Individual values are shown along with mean ± s.e.m. **E)** Same as (**D**) for optogenetically-evoked EPSCs. No significant differences were observed between groups (amplitude: p = 0.16; latency: p = p=0.15; rise time: p = 0.26; Kruskall-Wallis).

We next bred the *Aldh1a1* knockout allele into *Dat*^IRES-Cre^ knockin mice, which express Cre recombinase in DAergic neurons (Bäckman et al., 2006) to enable adeno-associated virus (AAV)-mediated expression of Cre-dependent channelrhodopsin 2 (ChR2) in SNc^DA^ neurons (Figures S1A and S1B). We obtained acute brain slices from mice in which Aldh1a1 expression is either intact (control) or knocked out, stimulated SNc^DA^ axons using brief flashes (1 ms) of full-field blue light (470 nm, 10 mW/mm^2^) and recorded GABA_A_ receptor-mediated inhibitory postsynaptic currents (IPSCs) from target SPNs in dorsal striatum using whole-cell voltage-clamp recordings (Figure 1 C). Consistent with previous reports (Kim et al., 2015), we found that the amplitude of optogenetically-evoked IPSCs (oIPSCs) was significantly reduced in slices obtained from *Aldh1a1*^-/-^ mice (0.71 ± 0.12 nA, n = 26 SPNs from 7 mice) compared to controls (1.25 ± 0.18 nA, n = 21 SPNs from 6 mice; Figure 1D). This effect could not be attributed to differences in our ability to stimulate SNc^DA^ axons with light, as optogenetically-evoked excitatory postsynaptic currents (oEPSCs) reflecting glutamate co-release were similar between conditions (Figures 1E and S1D). Importantly, large oIPSCs –including some exceeding 1 nA– were still detected in SPNs from *Aldh1a1*^-/-^ mice and were similar to controls in synaptic latency (control: 1.89 ± 0.09 ms; Aldh1a1^-/-^: 2.03 ± 0.10 ms; Figure 1D), rise time (control: 1.62 ± 0.08 ms; Aldh1a1^-/-^: 1.82 ± 0.11 ms; Figure 1D) and sensitivity to the GABA_A_ receptor antagonist SR95531 (Figure S1C), indicating that Aldh1a1 is not strictly required for SNc^DA^ neurons to synaptically inhibit SPNs.

Because Aldh1a1 expression is not exclusive to SNc (Figure 1A; Adam et al., 2012; Goto et al., 2018; Saunders et al., 2018; Zhang et al., 2014), we next sought to determine whether the reduction in GABA co-release observed in *Aldh1a1*^-/-^ mice results from the specific loss of Aldh1a1 in SNc^DA^ neurons. If Aldh1a1 functions as an intracellular GABA synthetic enzyme, we reasoned that selectively restoring its expression in SNc^DA^ neurons ought to rescue inhibitory co-transmission in *Aldh1a1*^-/-^ mice. We therefore generated an AAV expressing Cre-dependent Aldh1a1, which we co-injected along with Cre-dependent ChR2 in the midbrain of *Dat*^IRES-Cre/+^;*Aldh1a1*^-/-^ mice (Figures 1B and S1A). Although this manipulation reestablished Aldh1a1 protein expression in the soma, dendrites and axons of midbrain DA neurons (Figure 1B), it failed to restore the amplitude of oIPSCs in SPNs (0.67 ± 0.13 nA; n = 24 SPNs from 6 mice; Figure 1D) without affecting oEPSCs (Figures 1E and S1D). These results indicate that Aldh1a1 expression in SNc^DA^ neurons does not directly contribute to ability of these cells to inhibit SPNs, and suggest instead that Aldh1a1 modifies inhibitory co-transmission from SNc^DA^ axons non-cell autonomously.

### Selective deletion of Gat1 from SNc^DA^ neurons abolishes GABAergic co-transmission

SNc^DA^ neurons in mice do not express the canonical GABA synthetic enzymes Gad65 and Gad67 (Poulin et al., 2014; Tritsch et al., 2014; Kim et al., 2015; Saunders et al., 2018). If Aldh1a1 does not produce GABA either, how do SNc^DA^ neurons synaptically inhibit SPNs? We previously proposed that SNc^DA^ neurons acquire GABA presynaptically using the plasma membrane transporter Gat1 (Tritsch et al., 2014). To test this hypothesis directly, we generated mice in which Gat1 can be conditionally knocked out (cKO) by Cre recombinase (referred to as *Gat1*^fl^ mice; see methods and Figures 2A and S2A–D). To validate our genetic strategy, we first deleted Gat1 constitutively in all tissues by breeding *Gat1*^fl^ mice to a line expressing Cre in the germline (*CMV*^Cre^ mice, with offspring referred to as Gat1 cKO^germline^). As shown in Figure 2B, Gat1 protein was undetectable throughout the brain of Gat1 cKO^germline^ mice by immunofluorescence, confirming our ability to ablate Gat1 in a Cre-dependent fashion.

**Figure 2.**
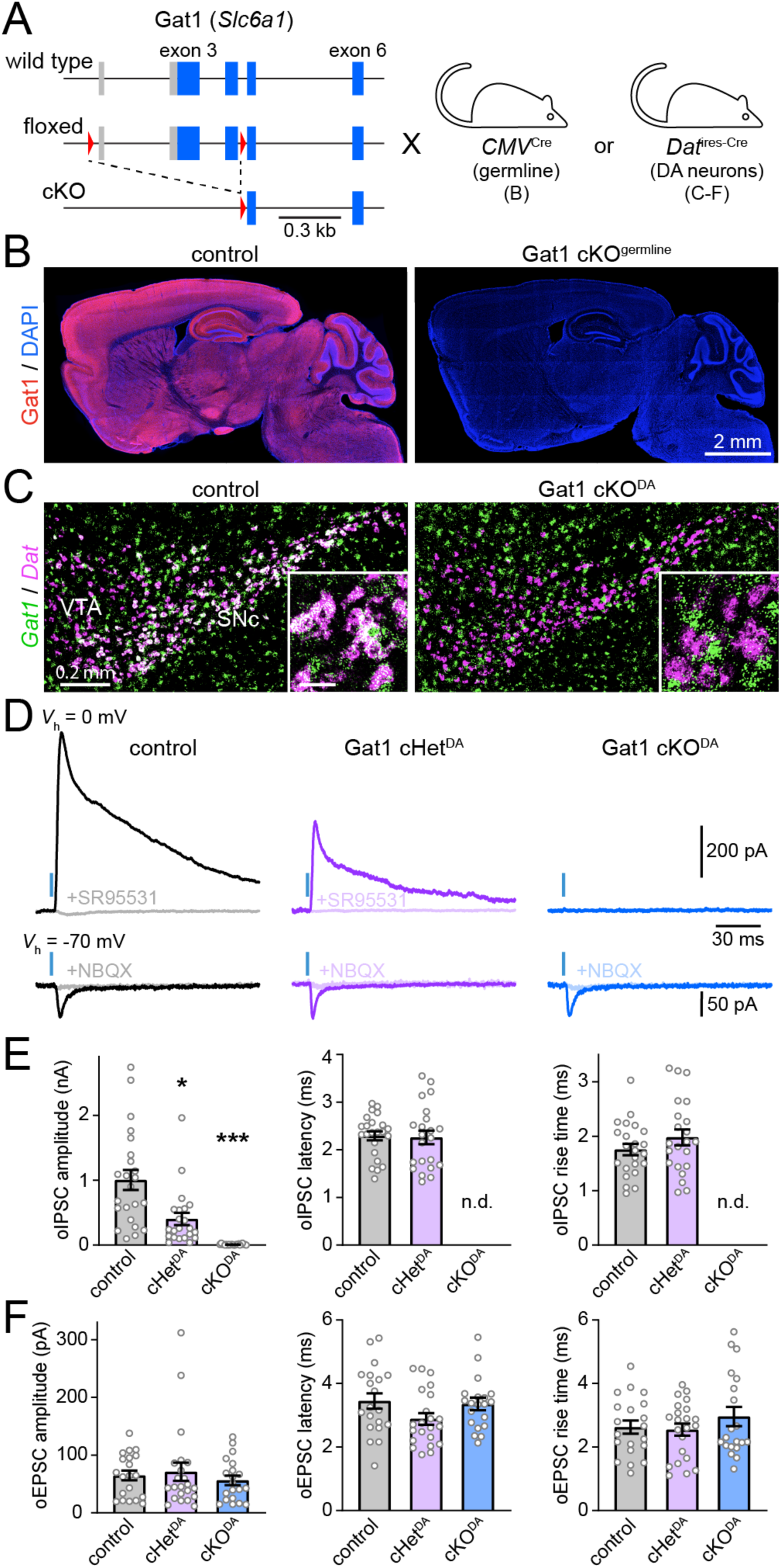
GABAergic co-transmission from SNc^DA^ neurons is abolished in Gat1 cKO^DA^ mice. **A)** Schematic of genetic strategy showing detail of Gat1’s endogenous locus (*Slc6a1*; top), the floxed allele generated by homologous recombination (see methods) before (middle) and after (bottom) Cre-mediated excision in *CMV*^Cre^ or *Dat*^IRES-Cre^ mice. Boxes represent exons with protein coding regions highlighted in blue, and red triangles depict LoxP sites. **B)** Sagittal brain sections from a control mouse (*Gat1*^+/+^; left) and a mouse homozygous for germline-deleted Gat1 (cKO^germline^; right) immunolabeled for Gat1 (red) and DAPI (blue). **C)** Epifluorescence image of coronal brain sections from a control mouse (*Dat*^IRES-Cre/+^;*Gat1*^+/+^; left) and a Gat1 cKO^DA^ mouse (*Dat*^IRES-Cre/+^;*Gat1*^fl/fl^; right) after two-color fluorescent *in situ* hybridization for *Gat1* (green) and *Dat* (magenta) to label midbrain DA neurons. Colocalization appears white. Inset: detail (scale bar: 50 µm). **D)** Inhibitory (top) and excitatory (bottom) postsynaptic currents recorded from SPNs held 0 and –70 mV, respectively, upon stimulation of ChR2-expressing nigrostriatal axons with blue light (1 ms, blue bar) in *Dat*^IRES-Cre/+^ mice with no (control; black), one (cHet^DA^; purple) or both (cKO^DA^; blue) alleles of Gat1 conditionally deleted from midbrain DA neurons before and after bath application of SR95531 (10 µM) or NBQX/CPP (10 µM). **E)** Quantification of optogenetically-evoked IPSC amplitude (left), synaptic latency (middle) and rise time (right) in slices from *Dat*^IRES-Cre/+^;*Gat1*^+/+^ (control), *Dat*^IRES-Cre/+^;*Gat1*^fl/+^ and *Dat*^IRES-Cre/+^;*Gat1*^fl/fl^ mice. Amplitude was strongly dependent on genotype (p = 2.3 × 10^-11^; Kruskall-Wallis; * p = 0.049 between control and Gat1 cHet^DA^, *** p = 1.2 × 10^-11^ between control and Gat1 cKO^DA^; Dunn’s Multiple Comparison). Latency: p = 0.67; rise time: p = 0.31 between control and Gat1 cHet^DA^; Mann–Whitney. n.d.: not detected. Individual values are show along with mean ± s.e.m. **F)** Same as (**E**) for optogenetically-evoked EPSCs. No significant differences were observed between groups (amplitude: p = 0.72; latency: p = p=0.10; rise time: p = 0.86; Kruskall-Wallis).

We next crossed *Gat1*^fl^ and *Dat*^IRES-Cre^ mice to generate offspring in which Gat1 is selectively deleted from DAergic cells (*Dat*^IRES-Cre/+^;*Gat1*^fl/fl^, henceforth referred to as Gat1 cKO^DA^ mice; Figures 2A, 2C and S2A). Using fluorescence *in situ* hybridization, we confirmed that Gat1 mRNA is no longer detected in SNc^DA^ neurons of Gat1 cKO^DA^ mice, yet is intact in surrounding inhibitory neurons within the substantia nigra (Figures 2C, S2E and S2F). To assess inhibitory co-transmission, we virally expressed Cre-dependent ChR2 in SNc^DA^ neurons and recorded light-evoked postsynaptic currents in SPNs in slices three or more weeks later. oIPSCs were readily observed in control slices (amplitude: 1.00 ± 0.15 nA; n = 23 SPNs from 8 mice), but were entirely absent in 22 SPNs from 6 Gat1 cKO^DA^ mice (Figures 2D, 2E and S2G). Importantly, the same SPNs voltage-clamped at the reversal potential for chloride showed oEPSCs that were similar to controls in prevalence, amplitude and kinetics (Figures 2D, 2F and S2H), confirming our ability to express ChR2 and optogenetically stimulate SNc^DA^ neurons in Gat1 cKO^DA^ mice. Moreover, we detected oIPSCs in mice heterozygous for the conditional allele of Gat1 (*Dat*^IRES-Cre/+^;*Gat1*^fl/+^ or Gat1 cHet^DA^ mice) that were 60% smaller than controls (0.40 ± 0.09 nA; n = 22 SPNs from 6 mice), but otherwise comparable in prevalence and kinetics (Figures 2D, 2E and S2G). These data suggest that Gat1 expression in SNc^DA^ neurons modulates the strength of inhibitory co-transmission onto SPNs in a dose-dependent fashion, with complete removal of Gat1 abolishing GABAergic signaling entirely.

### Deleting Gat1 from SNc^DA^ neurons does not impair basic nigrostriatal physiology

Preventing glutamate co-transmission from midbrain DA neurons impairs DA axon development and vesicular DA release (Hnasko et al., 2010; Fortin et al., 2012). It is therefore conceivable that our inability to evoke oIPSCs in SPNs following SNc^DA^ neuron stimulation in Gat1 cKO^DA^ mice results from non-specific effects of our genetic manipulation on nigrostriatal function. To test this possibility, we first determined whether loss of Gat1 affects the development of SNc^DA^ neurons using immunofluorescence for the DA synthetic enzyme tyrosine hydroxylase (TH). Figures 3A–3C show that the density of TH-positive cell bodies in midbrain and axons in dorsal striatum were indistinguishable in control and Gat1 cKO^DA^ mice. We also did not observe any difference in TH or Aldh1a1 protein levels, or in the cellular distribution of Aldh1a1 in SNc^DA^ neurons across genotypes (Figures 3D and 3E), indicating that deleting Gat1 from midbrain DA neurons does not grossly impair their development.

**Figure 3.**
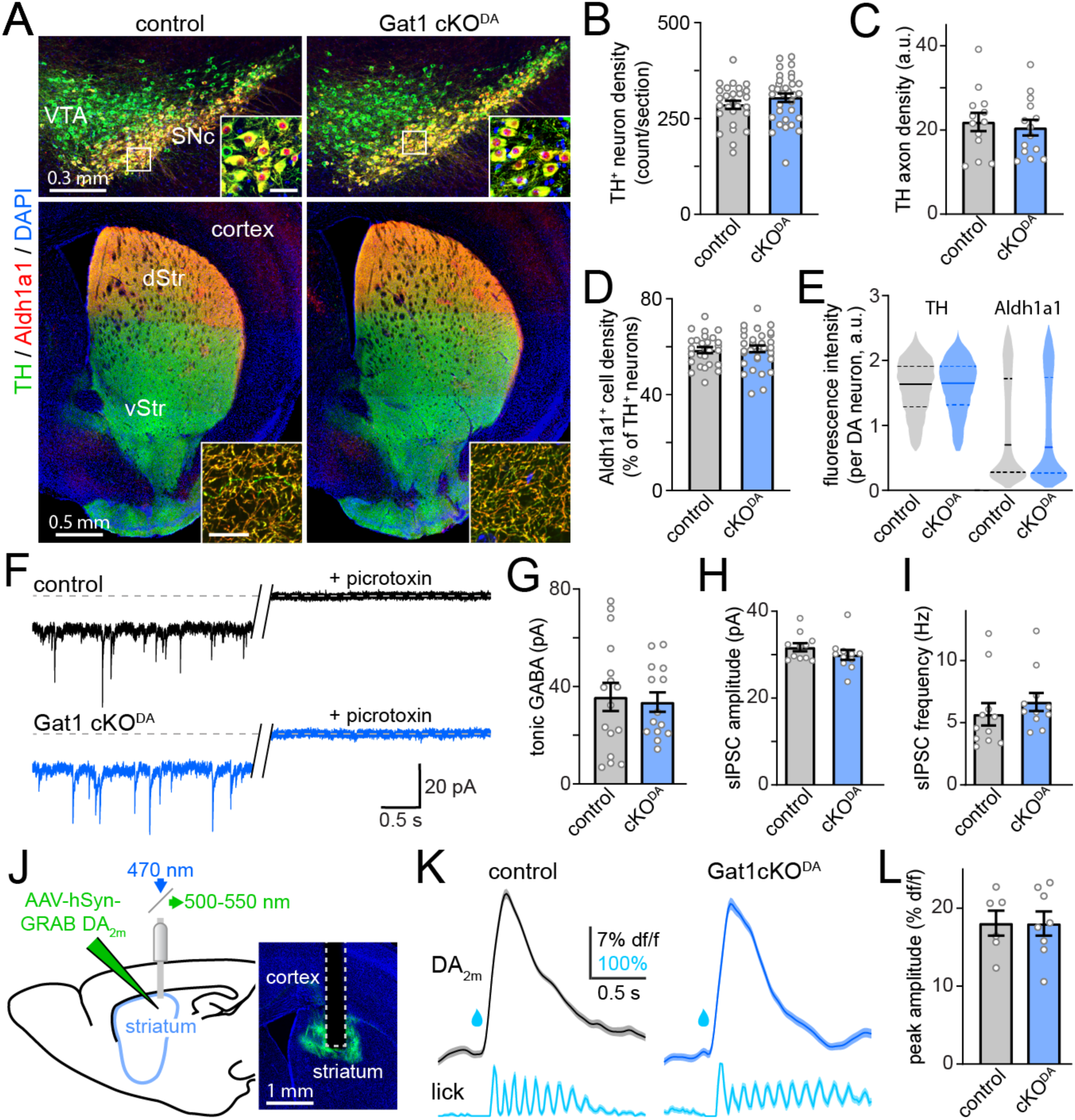
Nigrostriatal development and function are largely intact in Gat1 cKO^DA^ mice. **A)** Coronal brain sections from control *Dat*^IRES-Cre/+^ (left) and *Dat*^IRES-Cre/+^;Gat1^fl/fl^ mouse (Gat1 cKO^DA^; right) mice immunolabeled for tyrosine hydroxylase (TH, green), Aldh1a1 (red) and DAPI (blue). Top row: confocal images of ventral midbrain with white box show magnified in inset (scale bar inset: 25 µm). Bottom row: Epifluorescence image of striatum. Inset: confocal image of dorsal striatum (scale bar: 20 µm). **B)** Total number of TH^+^ DA neurons identified per midbrain section in control (n = 27 sections from 4 mice) and Gat1 cKO^DA^ mice (n = 32 sections from 5 mice; p = 0.18 vs. control, Mann– Whitney ). **C)** Mean density of TH^+^ axons imaged in dorsal striatum of control (n = 13 sections from 3 mice) and Gat1 cKO^DA^ mice (n = 14 sections from 3 mice; p = 0.76 vs. control, Mann–Whitney). **D)** Same as (**B**) for fraction of TH^+^ DA neurons positive for Aldh1a1 (p = 0.67, Mann–Whitney). **E)** Violin plot of TH and Aldh1a1 relative fluorescence intensity per TH^+^ DA neuron in control (n = 7714 cells from 4 mice) and Gat1 cKO^DA^ mice (n = 9740 cells from 5 mice). Filled line indicates median, and dashed lines first and third quartiles. **F)** Example whole-cell voltage-clamp recordings (*V*_h_ = –70 mV) from SPNs in control (black) and Gat1 cKO^DA^ slices (blue) before and after the application of the GABA_A_ receptor blocker picrotoxin (100 µM). Slices were continuously bathed in NBQX (10 µM), R-CPP (10 µM), and CGP55845 (5 µM) to block ionotropic glutamate and metabotropic GABA_B_ receptors. The shift in picrotoxin-evoked holding current highlights the inward tonic GABAergic current at baseline. **G)** Quantification of picrotoxin-evoked shift in holding currents recorded in control (n = 16 SPNs from 6 mice) and Gat1 cKO^DA^ slices (n = 14 SPNs from 5 mice; p = 0.92 vs. control, Mann– Whitney). **H)** Mean amplitude of spontaneous IPSCs recorded from SPNs in control (n = 11 SPNs from 4 mice) and Gat1 cKO^DA^ slices (n = 11 SPNs from 4 mice; p = 0.24 vs. control, Mann–Whitney). **I)** same as (**H**) for sIPSC frequency (p = 0.10, Mann–Whitney). **J)** Left: schematic of photometry experiment. Right: example coronal section showing fiber tract in dorsal striatum expressing the fluorescent DA sensor GRAB-DA_2m_. **K)** Mean (± s.e.m.) GRAB-DA_2m_ fluorescence (top) and lick probability (bottom) following water delivery (drop) in control (left) and Gat1 cKO^DA^ mice (right). **L)** Mean reward-evoked GRAB-DA_2m_ peak fluorescence in control (n = 6 imaging sessions from 3 mice) and Gat1 cKO^DA^ mice (n = 8 sessions from 4 mice; p = 0.95 vs. control, Mann–Whitney). Data in **B-E**, **G-I** and **L** show Individual values along with mean ± s.e.m.

Gat1 plays an important role in limiting the buildup of extracellular GABA in striatum (Ade et al., 2008; Kirmse et al., 2008; Tritsch et al., 2014; Roberts et al., 2020). Recent studies show that elevating extracellular GABA significantly dampens the excitability of SNc^DA^ axons through activation of presynaptic GABA_A_ and GABA_B_ receptors (Brodnik et al., 2019; Lopes et al., 2019; Roberts et al., 2020; Kramer et al., 2020). Deleting Gat1 from SNc^DA^ neurons may therefore depress their ability to release transmitters indirectly by decreasing GABA reuptake capacity in striatum and causing ambient GABA levels to rise. To test this, we estimated extracellular GABA levels in striatum by measuring the amplitude of tonic GABA_A_ receptor-mediated currents in SPNs with the GABA_A_ receptor pore blocker picrotoxin. Picrotoxin-mediated shifts in holding current were not different in control and Gat1 cKO^DA^ slices (Figures 3F and 3G), suggesting that extracellular GABA levels are comparable between conditions.

We also found no differences in the membrane resistance or capacitance of SPNs (Figures S2I and S2J), or in the amplitude and frequency of spontaneous IPSCs recorded from SPNs (Figures 3H and 3I). These results indicate that the membrane properties of SPNs and their capacity to detect ambient and synaptic GABAergic currents are not affected in Gat1 cKO^DA^ mice. Postsynaptic deficiencies are therefore unlikely to account for our failure to detect GABAergic responses from SNc^DA^ neurons.

Lastly, we assessed the integrity of the vesicular release machinery of SNc^DA^ neurons in Gat1 cKO^DA^ mice. Although glutamatergic co-transmission from SNc^DA^ neurons appears intact (Figure 2F), glutamate release is anatomically and molecularly distinct from DA’s (Fortin et al., 2019; Silm et al., 2019; Zhang et al., 2015). By contrast, DAergic and GABAergic transmission from SNc^DA^ neurons both depend on the vesicular monoamine transporter Vmat2 and presumably occur from the same presynaptic specializations (Tritsch et al., 2012; Stensrud et al., 2014). We therefore compared the magnitude of DA transients evoked by water delivery in water-restricted control and Gat1 cKO^DA^ mice using fiber photometry of the fluorescent DA sensor GRAB-DA_2m_ in striatum (Figures 3J; Sun et al., 2020). As shown in Figures 3K and 3L, phasic DA transients evoked during water consumption were not smaller in Gat1 cKO^DA^ mice compared to controls, indicating that activity-dependent DA exocytosis is not compromised in SNc^DA^ neurons lacking Gat1. Together, these results point to Gat1 cKO^DA^ mice having a selective deficit in the ability of SNc^DA^ neurons to liberate the transmitter that inhibits SPNs.

### Inhibitory co-transmission relies on Gat1-mediated import of GABA

How does Gat1 contribute to GABAergic signaling from SNc^DA^ neurons? Gat1 normally transports GABA from the synaptic cleft into the cytoplasm using energy from the Na^+^ and Cl^-^ electrochemical gradients across the plasma membrane (Scimemi, 2014). In SNc^DA^ neurons, Gat1 may therefore serve as a source of cytosolic GABA for vesicular loading and synaptic release. Alternatively, Gat1 may operate in reverse during neuronal depolarization to mediate non-vesicular GABA efflux (Wu et al., 2007; Bertram et al., 2011), leaving the source of GABA in SNc^DA^ neurons to be identified. Lastly, Gat1 may be part of a molecular complex that controls inhibitory signaling from SNc^DA^ neurons independently of its transporter function (Deken et al., 2000; Ryan et al., 2021). To distinguish these possibilities, we attempted to restore inhibitory transmission in Gat1 cKO^DA^ mice by selectively expressing constructs in SNc^DA^ neurons.

We first made a Cre-dependent AAV encoding wild-type Gat1 tagged with influenza hemagglutinin (HA) at its C-terminus, which we injected in the midbrain of adult Gat1 cKO^DA^ mice along with Cre-dependent-ChR2. Immunostaining for HA confirmed our ability to reintroduce Gat1 specifically in SNc^DA^ neurons (Figure 4A). We observed Gat1-HA throughout the axons of SNc^DA^ neurons in dorsal striatum (Figure S3A), consistent with Gat1’s usual presynaptic distribution (Scimemi, 2014; Uchigashima et al., 2016; Fattorini et al., 2020). Importantly, exogenous expression of Gat1-HA in SNc^DA^ neurons of Gat1 cKO^DA^ mice fully restored GABAergic signaling (Figures 4C and 4H–J), as oIPSCs were indistinguishable from those recorded in wild type controls in amplitude (1.49 ± 0.23 nA; n = 14 SPNs from 4 mice), synaptic latency (2.25 ± 0.18 ms) and rise time (2.14 ± 0.24 ms; all p > 0.05 vs. control, Mann– Whitney). Together, these data show that SNc^DA^ neurons remain capable of inhibiting SPNs in adult Gat1 cKO^DA^ mice if provided with Gat1.

**Figure 4.**
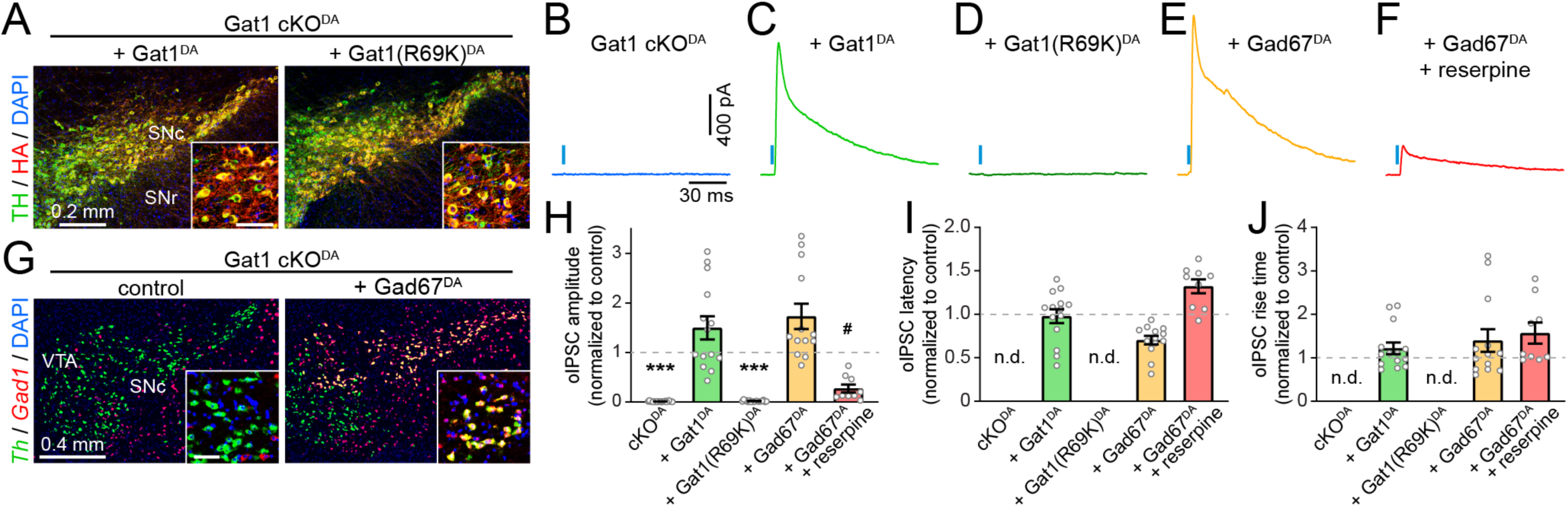
GABA co-release from SNc^DA^ axons requires Gat1-mediated GABA uptake. **A)** Confocal images of coronal ventral midbrain sections from *Dat*^IRES-Cre/+^;*Gat1*^fl/fl^ mice virally expressing Cre-dependent HA-tagged Gat1 (left) and Gat1(R69K) (right) in SNc immunolabeled for TH (green), HA (red) and DAPI (blue). White box magnified in inset (scale bar: 50 µm). **B–E)** Example IPSCs recorded from SPNs (*V*_h_ = 0 mV) upon stimulation of ChR2-expressing SNc^DA^ axons (1 ms, blue bar) in Gat1 cKO^DA^ mice (**B**) exogenously expressing Gat1 (**C**), Gat1(R69K) (**D**) or Gad67 (**E**). **F)** Same as (**E**) after treatment with reserpine. **G)** Two-color fluorescent *in situ* hybridization for Gad67 (red) and tyrosine hydroxylase (green) in Gat1 cKO^DA^ mice with (right) or without (left) AAV-mediated expression of Gad67 in DA neurons. Inset: high magnification detail (scale bar: 50 µm). **H)** Quantification of oIPSC amplitude in SPNs of Gat1 cKO^DA^ mice virally expressing in SNc^DA^ neurons ChR2 only (n = 22 SPNs from 6 mice, p = 4.9 × 10^-13^), or ChR2+Gat1 (n = 14 SPNs from 4 mice; p = 0.08), ChR2+Gat1(R69K) (n = 18 SPNs from 5 mice; p = 9.9 × 10^-12^), ChR2+Gad67 (n = 13 SPNs from 4 mice; p = 0.047) and ChR2+Gad67 following reserpine treatment (n = 9 SPNs from 2 mice; p = 4 × 10^-6^ vs. ChR2+Gad67) normalized to responses in control mice (*Dat*^IRES-Cre/+^;*Gat1*^+/+^). All p-values vs. control unless specified otherwise, Mann-Whitney. n.d.: not detected. **I)** same as **(H)** for latency (Gat1^DA^: p= 0.98 vs. control; Gad67^DA^: p = 0.0002 vs. control; Gad67^DA^+reserpine: p = 8 × 10^-6^ vs. Gad67, Mann-Whitney). **J)** same as **(I)** for rise time (Gat1^DA^: p = 0.32 vs. control; Gad67^DA^: p = 0.16 vs. control; Gad67^DA^+reserpine: p = 0.44 vs. Gad67; Mann–Whitney).

To distinguish whether the dependence on Gat1 stems from its ability to transport GABA across the plasma membrane or interact with other molecules, we made another Cre-dependent AAV encoding Gat1-HA with a single amino acid mutation (R69K) that abolishes transmembrane GABA transport but not membrane expression (Dayan-Alon and Kanner, 2019; Pantanowitz et al., 1993). We co-expressed this construct along with ChR2 in midbrain DA neurons of Gat1 cKO^DA^ mice and assessed inhibitory co-transmission onto SPNs. Despite strong expression of Gat1(R69K)-HA in TH-positive SNc^DA^ cell bodies and axons (Figures 4A and S3A), and successful ChR2-mediated activation of SNc^DA^ axons in striatum (Figures S3B–E), we were unable to evoke oIPSCs in 18 SPNs from 5 mice (Figures 4D and 4H). This suggests that SNc^DA^ neurons depend on Gat1’s transport function to inhibit SPNs.

If Gat1 acts presynaptically to import GABA for vesicular release, a strong prediction is that other means of supplying cytosolic GABA to SNc^DA^ axons of Gat1 cKO^DA^ mice should reinstate inhibitory transmission. To test this, we virally transduced SNc^DA^ neurons of Gat1 cKO^DA^ mice with ChR2 and Gad67, which catalyzes the decarboxylation of glutamate to yield GABA throughout the cytosol of GABAergic neurons (Asada et al., 1997; Martin and Rimvall, 1993). Consistent with our prediction, expressing Gad67 in SNc^DA^ neurons lacking Gat1 (Figure 4G) rescued inhibitory co-transmission (Figures 4E and 4H–J), yielding oIPSCs that were slightly larger in amplitude (1.72 ± 0.26 nA; n = 13 SPNs from 4 mice; p = 0.047 vs. control, Mann– Whitney) and faster in onset than controls (1.61 ± 0.11 ms; p = 0.0002 vs. control, Mann– Whitney). This shows that Gad67 can effectively substitute for Gat1 to sustain inhibitory transmission from SNc^DA^ neurons. We obtained similar results with another GABA synthetic enzyme (Gad65; Figures S3F and S3G) that preferentially associates with synaptic vesicles in axonal terminals (Kaufman et al., 1991; Tian et al., 1999). Interestingly, oIPSCs evoked from SNc^DA^ neurons expressing Gad65 or Gad67 were significantly less prone to rundown with successive optogenetic stimulation compared to Gat1-expressing controls (Figure S3H). Together, these experiments show that Gat1 contributes to inhibitory co-transmission by importing GABA into the cytoplasm of SNc^DA^ neurons for vesicular loading.

We previously suggested that GABAergic transmission from SNc^DA^ neurons requires the vesicular monoamine transporter Vmat2 (Tritsch et al., 2012). To confirm that Vmat2 contributes to vesicular packaging of GABA, we treated slices from Gat1 cKO^DA^ mice expressing Gad67 in SNc^DA^ neurons with the selective Vmat2 antagonist reserpine. Reserpine treatment strongly depressed the amplitude of light-evoked IPSCs (0.27 ± 0.08 nA; n = 9 SPNs from 2 mice; p = 4 × 10^-6^ vs. Gat1 cKO^DA^ + Gad67^DA^, Mann–Whitney ; Figures 4F and 4H), but not EPSCs (Figures S4B–E), providing strong evidence that Vmat2 supports vesicular transport of GABA. Together, these data demonstrate that the neurotransmitter that SNc^DA^ neurons release is GABA, and that Gat1 acts presynaptically in SNc^DA^ neurons to supply cytosolic GABA for Vmat2-dependent vesicular release.

## DISCUSSION

SNc^DA^ neurons are essential for learning and producing goal-directed movements, but the mechanisms through which they impact the activity of target cells to affect behavior are still poorly understood. In addition to releasing DA, SNc^DA^ neurons exert a rapid and potent inhibitory influence on downstream striatal neurons through synaptic co-release of an inhibitory transmitter, the identity of which has remained speculative (Tritsch et al., 2016). Here, we show that the co-released transmitter is GABA, that vesicular co-release of GABA depends on Vmat2, and that SNc^DA^ neurons sustain inhibitory transmission without producing GABA, relying instead on presynaptic uptake through the neuronal plasma membrane GABA transporter Gat1.

The chemical identity of a synaptic neurotransmitter is traditionally established by determining if a synapse possesses the means to (a) acquire said transmitter presynaptically, typically by expressing synthetic enzymes, (b) package it into synaptic vesicles, (c) detect it postsynaptically using specific receptors, and (d) terminate its synaptic actions, most commonly through membrane uptake or enzymatic degradation. SNc^DA^ neuron stimulation evokes nanoampere-sized GABAA receptor-mediated IPSCs in SPNs (Ishikawa et al., 2013; Kim et al., 2015; Nelson et al., 2014; Tritsch et al., 2012), and the decay kinetics of IPSCs are shaped by membrane GABA transporters in striatum (Tritsch et al., 2014). These observations strongly suggest that the released transmitter is GABA. However, this assertion was called in question, as mouse SNc^DA^ neurons lack the molecular machinery considered essential for GABA release. Indeed, they do not express the GABA synthetic enzymes Gad65 or Gad67, or the vesicular GABA transporter Vgat (Kim et al., 2015; Poulin et al., 2014; Saunders et al., 2018; Tritsch et al., 2012, 2014). In addition, inhibitory transmission from SNc^DA^ neurons requires Vmat2 (Tritsch et al., 2012), which is not known to transport GABA (Yelin and Schuldiner, 1995), and GABA_A_ receptors can be activated by a variety of ligands and allosteric modulators in addition to GABA (Belelli and Lambert, 2005; Johnston, 1996).

In this study, we generate a conditional knockout of the membrane GABA transporter Gat1 and show that inhibitory co-transmission from SNc^DA^ neurons is entirely abolished upon selective deletion of Gat1 from DA neurons. We demonstrate that this effect is not secondary to developmental or experimental confounds in our conditional knockout, as SPNs maintain their ability to detect synaptically-released GABA, and SNc^DA^ neurons remain structurally and functionally competent to co-release DA, glutamate, and even GABA (if GABA is provided to DA neurons through cell type-specific expression of Gad65 or Gad67). Importantly, we restore inhibitory co-transmission from SNc^DA^ neurons in adult mice by selectively introducing a wild-type allele of Gat1 in midbrain DA neurons, but not a GABA transport-deficient point-mutant. These findings therefore indicate that Gat1’s transporter function is required for SNc^DA^ neurons to synaptically inhibit SPNs, and that Gad65 and Gad67 can functionally replace Gat1. When combined with previous reports showing that GABA can be observed in close association with Vmat2-containing synaptic vesicles in striatal DA terminals (Stensrud et al., 2014) and that Vgat can substitute for Vmat2 to sustain inhibitory transmission from SNc^DA^ neurons (Tritsch et al., 2012), our results firmly establish GABA as the transmitter that SNc^DA^ neurons release along with DA and glutamate.

More generally, our results provide important mechanistic insights into GABAergic synaptic transmission. First, our study describes the first instance of a mammalian GABAergic synapse lacking the ability to produce GABA. In the central nervous system of mice, GABA is almost exclusively synthetized *de novo* from glutamate by Gad65 and Gad67, with Gad67 accounting for most of the GABA produced (Asada et al., 1996, 1997; Tian et al., 1999; Martin and Rimvall, 1993). Gad65 and Gad67 are therefore considered critical elements for conferring neurons a GABAergic identity. However, our results call for an expansion of this list, given our demonstration that SNc^DA^ neurons rely exclusively on Gat1-mediated presynaptic uptake as their source of GABA. The complete reliance of SNc^DA^ axons on GABA uptake raises the interesting possibility GABA co-release may locally vary at individual synapses with extracellular GABA levels and Gat1 transporter activity. More importantly, our findings suggest that the function of presynaptic plasma membrane GABA transporters extends beyond terminating GABA’s actions in the synaptic cleft to include sustaining vesicular filling. Such presynaptic contribution has proven difficult to dissect at Gad-containing synapses (Bragina et al., 2008; Jensen et al., 2003; Wang et al., 2013). The ability of Gat1 to sustain GABA release *in vivo* is not known, but likely exceeds that in brain slices, as extracellular GABA levels *ex vivo* are reduced by the absence of action potential-dependent GABA exocytosis in striatum and by the dilution of released GABA in the perfusate. Indeed, synaptic release of DA – which is similarly dependent on presynaptic uptake by the membrane DA transporter Dat – also strongly depresses during repeated stimulation in brain slices (Condon et al., 2019; Liu et al., 2021; O’Neill et al., 2017; Sulzer et al., 2016), yet voltammetry and photometry recordings show sustained release *in vivo* (Patriarchi et al., 2018; Stamford et al., 1987; Sun et al., 2020; Wightman and Zimmerman, 1990).

Second, they reveal that, in the absence of Gat1, SNc^DA^ neurons do not possess GABA for synaptic release. This is at odds with a recent report claiming that Aldh1a1 produces GABA *de novo* in SNc^DA^ neurons, accounting for over half of co-released GABA (Kim et al., 2015). This finding helped establish aldehyde dehydrogenases as GABA synthetic enzymes in the central nervous system of mammals, significantly expanding the number of cells capable of influencing neural function through GABAergic signaling (Kwak et al., 2020; Jin et al., 2021; Melani and Tritsch, 2021). Yet, the pharmacological and genetic manipulations employed by Kim and colleagues to establish GABA as a product of Aldh1a1 activity lacked specificity for SNc^DA^ neurons. Here, we independently replicate the observation that GABA co-release is significantly diminished in mice lacking Aldh1a1 in all cells throughout development. However, we were unable to restore inhibitory co-transmission by selectively re-introducing Aldh1a1 in SNc^DA^ neurons (Figure 1). By contrast, selectively deleting Gat1 from DA neurons completely abolished GABA co-release (Figure 2) without altering Aldh1a1 expression (Figures 3D and 3E). Moreover, IPSCs continue to be detected from SPNs in Gat cKO^DA^ mice if SNc^DA^ neurons are provided with a synthetic enzyme (Figures 4E–H). Together, these data show that Aldh1a1 does not directly provide GABA to SNc^DA^ neurons, and suggest instead that Aldh1a1 modifies GABA co-transmission non-cell autonomously. Aldh1a1 is widely expressed throughout the nervous system of humans and mice (Adam et al., 2012; Alnouti and Klaassen, 2008; Goto et al., 2018; Saunders et al., 2018; Zhang et al., 2014), where it controls numerous biological processes that may impact GABAergic signaling indirectly. For instance, Aldh1a1 is one of three enzymes responsible for converting retinal into retinoic acid, an important developmental morphogen and transcriptional regulator (Duester, 2008; Marchitti et al., 2008; Rhinn and Dollé, 2012). Interfering with Aldh1a1 disrupts the development of DA axons and mu opioid receptor-rich patches in striatum (Jacobs et al., 2007; Sgobio et al., 2017; Pan et al., 2019). Aldh1a1 also maintains cellular homeostasis by degrading reactive oxygen species and toxic acetaldehyde metabolites, and by limiting the aggregation of misfolded proteins. In DA neurons in particular, loss of Aldh1a1 leads to the accumulation of toxic DA metabolites and α-synuclein aggregates that may affect GABA release (Anderson et al., 2011; Burke et al., 2008; Goldstein et al., 2013; Liu et al., 2014; Marchitti et al., 2007; Roberts et al., 2020). Lastly, aldehyde dehydrogenases in astrocytes modulate synaptic transmission by altering extracellular GABA homeostasis and neuronal excitability (Kwak et al., 2020; Jin et al., 2021; Melani and Tritsch, 2021).

Lastly, although Vgat is typically considered essential for packaging GABA into synaptic vesicles, Vgat knockout mice are incompletely deficient in fast synaptic release of GABA (Wojcik et al., 2006), suggesting that alternative vesicular GABA transporters exist. Our results provide evidence that Vmat2 directly contributes to vesicular packaging of GABA along with DA as reserpine blocks IPSCs evoked from Gat1 cKO^DA^ mice expressing Gad67. The finding further alleviates concerns that Vmat2 antagonists indirectly affects GABAergic transmission from SNc^DA^ axons by non-specifically compromising Gat1 function. Because GABA lacks the characteristic charge and structure of established Vmat2 substrates (Yelin and Schuldiner, 1995), our findings indicate that the biology of this important transporter remains incompletely understood. Future investigations will determine whether Vmat2 interacts with other vesicular proteins to load GABA into synaptic vesicles and whether DA and GABA are co-packaged into the same vesicles or released from distinct presynaptic specializations. Together, our data redefine the minimal molecular machinery necessary to impart neurons with a GABAergic phenotype and pave the way for studies dissecting the specific contribution of GABA co-release from SNc^DA^ neurons to behavior. In addition, our work provides a novel genetic tool with which to dissect the cell-type specific contribution of Gat1 to neural function and dysfunction in Parkinson’s disease, epilepsy and autism-spectrum disorders (Kahen et al., 2021; Madsen et al., 2010; Roberts et al., 2020; Sanders et al., 2015).

## ACKNOWLEDGEMENTS

We thank Adam Carter, Margaret Rice, Dick Tsien and members of the Tritsch laboratory for comments on the manuscript, Yulong Li for the GRAB-DA_2m_ DA sensor, and Huaibin Cai for kindly providing Aldh1a1^-/-^ mice. We acknowledge the New York University Medical Center Rodent Genetic Engineering Laboratory for rederivation, the Genotyping Core Laboratory for mouse genotyping, the Department of Comparative Medicine for animal care and maintenance and the Neuroscience Institute’s imaging facilities for microscopes availability. R.M. is supported by a Marlene and Paolo Fresco Postdoctoral Fellowship. N.X.T. is an Alfred P. Sloan Research Fellow in Neuroscience, and is supported by grants from the Dana, Whitehall and Feldstein Medical Foundations, as well as from the National Institutes of Health (DP2NS105553).

## AUTHOR CONTRIBUTIONS

R.M. and N.X.T. designed and performed experiments, analyzed and interpreted the data, and wrote the manuscript.

## DECLARATION OF INTERESTS

The authors declare no competing interests.

## METHODS

### Experimental animals

Adult mice (8-28 weeks old) of both sexes were used in this study in accordance with protocols approved by the New York University Langone Health Institutional Animal Care and Use Committee (protocol #170123). Mice were housed in groups under a reverse 12 hr light-dark cycle (dark from 10 a.m. to 10 p.m.) with ad libitum access to food and water. Knock-in mice bearing an internal ribosome entry site (IRES)-linked Cre recombinase gene downstream of the gene *Slc6a3*, which encodes the plasma membrane DA transporter Dat (referred to as *Dat*^IRES−Cre^ mice) were obtained from the Jackson Laboratory (strain # 006660), along with CMV^Cre^ mice (strain # 006054). Mice in which the gene encoding Aldh1a1 is conventionally knocked out (B6.129-*Aldh1a1^tm1Gdu^*/J; referred to as Aldh1a1^-/-^ mice) were kindly donated by Dr. Huaibin Cai (NIH). Transgenic lines were maintained on a C57Bl6/J background (Jackson Laboratory strain # 000664) and bred to one another to generate experimental animals, which were genotyped both in house by the NYU Genotyping Core and by Transnetyx.

## Generation of conditional Gat1 knockout mice

Gat1^fl^ mice were generated by genOway by homologous recombination in mouse embryonic stem (ES) cells. LoxP sites flank exons 2, 3 and 4 of *Slc6a1* (Figures 2A and S2A), which contain the ATG initiation codon, the N-terminal intracellular tail, 2 sodium binding sites essential for GABA transport and 3 of 12 transmembrane domains (Bendahan and Kanner, 1993; Pantanowitz et al., 1993). The targeting vector contained a cassette encoding diphteria toxin and a neomycin resistance cassette flanked by FRT sites for negative and positive selection of ES cell clones, respectively. Homologous recombination at the intended locus were confirmed by PCR screening and Southern blotting of ES cell clones’ genomic DNA (Figure S2B). ES cell clones positive for the knock-in (ki) allele were injected into blastocysts to obtain chimeric males, which were bred to Flp deleter mice to excise the Neomycin selection cassette and generate heterozygous mice carrying the floxed allele of Gat1 (referred to as Gat1^fl^ mice). Crossing these mice with lines expressing Cre recombinase enabled the excision of the loxP-flanked region, resulting in a Gat1 conditional knockout allele (cKO): *CMV*^Cre^ mice mediate germline recombination to knock Gat1 out throughout the body, while *Dat*^IRES-Cre^ mice delete postnatally Gat1 from Dat-expressing DA neurons. The following primers were designed and validated by GenOway for the specific detection Gat1^fl^ and Gat1^KO^ alleles: GAT1-FloxF and GAT1-FloxR yielding 170 bp wild-type and 276 bp conditional knock-out bands (Figure S2C), and GAT1-Flox14F, GAT1-Flox18F and GAT1-Flox15R yielding 211 bp wild-type and 818 bp constitutive knock-out bands (Figure S2D).

## Adeno-associated viruses (AAVs)

Cre-dependent AAV viruses were used to selectively transduce midbrain DA neurons with channelrhodopsin 2 (ChR2) and express constructs to rescue GABA co-release in *Dat*^IRES-Cre/+^;*Aldh1a1*^-/-^ and *Dat*^IRES-Cre/+^;*Gat1*^fl/fl^ (Gat1 cKO^DA^) mice. AAV8-EF1a-double floxed-hChR2(H134R)-mCherry-WPRE-HGHpA was obtained from Addgene (# 20297) and AAV9-hSyn-GRAB_DA2m was provided by Dr. Yulong Li. The following AAVs were generated by replacing the coding sequence of ChR2-mCherry in Addgene’s plasmid #20297 with the protein-coding sequence for Aldh1a1 (NM_013467.3), Gat1 (NM_178703.4), Gat1(R69K), Gad65 (NM_008078.2) or Gad67 (NM_008077.5) using Asc1 and Nhe1 restriction sites (Genscript). Three copies of the influenza hemagglutinin (HA) coding sequence were inserted at the C-terminus of all constructs except for Gad65 and Gad67 to facilitate immunohistochemical identification. Viral vectors were subsequently packaged (serotype 8) by the Boston Children Hospital viral core, stored in undiluted aliquots at −80°C and diluted 2-10X in sterile 0.9% saline immediately prior to intracranial injection.

## Stereotaxic surgery

For stereotaxic viral injections, mice were anesthetized with isoflurane, placed in an animal stereotaxic frame (David Kopf Instruments) and injected (rate: 100 nl/min) in the right SNc of *Dat*^IRES-Cre^ mice with 1 μl of a mix of AAVs encoding Cre-dependent ChR2 and either Cre-dependent Aldh1a1, Gat1, Gat1(R69K), Gad65, Gad67 or saline using a syringe pump (Legato 111; KD scientific) fitted with a Hamilton syringe (Gastight 1701N) connected to a pulled glass injection micripipet (∼100 μm tip; Wiretrol II; Drummond) via PE tubing filled with mineral oil. The optimal dilution of each AAV was empirically determined to enable strong expression while avoiding toxicity. Injection coordinates for SNc were (from bregma): AP –3.12 mm, ML +1.6 mm and DV –4.2 mm (from pia). For fiber photometry, 250 nl of AAV9-hSyn-GRAB_DA_2m_ was injected 2.2 mm below pia into the right dorsal striatum (from bregma: AP, +0.5mm; ML, +2.0 mm). Mice were allowed to recover for at least 3 weeks prior to electrophysiology.

## Slice electrophysiology

Acute brain slices and whole-cell voltage-clamp recordings from SPNs were obtained using standard methods, as described previously (Tritsch et al., 2012, 2014). Briefly, mice were anesthetized and perfused with ice-cold artificial cerebrospinal fluid (ACSF) containing (in mM) 125 NaCl, 2.5 KCl, 25 NaHCO3, 2 CaCl2, 1 MgCl2, 1.25 NaH2PO4 and 11 glucose (295 mOsm·kg−1). Sagittal slices of striatum (275-μm thick) were subsequently obtained in cold choline-based cutting solution (in mM: 110 choline chloride, 25 NaHCO3, 2.5 KCl, 7 MgCl2, 0.5 CaCl2, 1.25 NaH2PO4, 25 glucose, 11.6 ascorbic acid, and 3.1 pyruvic acid) using a Leica VT1200 S vibratome. Following 15 min recovery in ACSF at 34°C, slices were kept at room temperature (20–22°C) until use. All solutions were constantly bubbled with 95% O2/5% CO2. For recording, slices were transferred to a Plexiglas chamber mounted on an upright microscope (SliceScope Pro 6000; Scientifica) and imaged through a 40X water-immersion objective (Olympus) with infrared light (780 nm) and differential interference contrast (DIC) optics using a complementary metal-oxide-semiconductor (CMOS) camera (ORCA-spark; Hamamatsu). Slices were held down by a flattened U-shaped platinum wire strung with individual strands of synthetic dental floss fibers and were continually superfused (2–2.5 ml/min) with ACSF at 33– 34°C by passing it through a feedback-controlled in-line heater (SH-27B; Warner Instruments) before entering the chamber. Whole-cell voltage-clamp recordings were established from SPNs in dorsal striatum identified visually and electrophysiologically. For oIPSC/oEPSC recordings, patch pipettes (2–3 MΩ) were filled with a cesium-based low-chloride internal solution containing (in mM): 135 CsMeSO_3_, 10 HEPES, 1 EGTA, 3.3 QX-314 (Cl_2_ salt), 4 Mg-ATP, 0.3 Na-GTP, 8 Na_2_-phosphocreatine (pH 7.3 adjusted with CsOH; 295 mOsm). Under these conditions, GABA_A_ receptor-mediated currents can be isolated as outward currents when SPNs are held at 0 mV, while glutamatergic currents can be isolated as inward currents at *E*_Cl_ = –70 mV. Inward tonic GABAergic currents and sIPSC in Figure 3 were recorded at -70 mV using an internal solution containing (in mM) 125 CsCl, 10 TEA-Cl, 10 HEPES, 0.1 Cs-EGTA, 3.3 QX-314 (Cl−salt), 4 Mg-ATP, 0.3 Na-GTP, 8 Na_2_-Phosphocreatine (pH 7.3 adjusted with CsOH; 295 mOsm) in the presence of NBQX, R-CPP, and CGP55845. For all voltage-clamp experiments, errors due to the voltage drop across the series resistance (<20 MΩ) were left uncompensated. Membrane potentials were corrected for a ∼5 mV liquid junction potential. To activate ChR2-expressing fibers, full-field 1 ms-long pulses of light from a 473-nm LED (p300 CoolLED) were delivered through the objective (20 mW mm^2^ under the objective) at 30 seconds intervals.

## Pharmacological reagents

Drugs (all from Tocris) were applied by bath perfusion: SR95531 (10 µM), picrotoxin (100 µM), CGP55845 (5 µM), 2,3-dihydroxy-6-nitro-7-sulfamoylbenzo(f)quinoxaline (NBQX; 10 µM), R,S-3-(2-carboxypiperazin-4-yl)propyl-1-phosphonic acid (R-CPP; 10 µM). To inhibit monoaminergic vesicular transport and deplete transmitter-filled vesicles, mice were injected intraperitoneal with the irreversible Vmat inhibitor reserpine (5 mg/kg) 24 h and 2 h before slicing. Brain sections from these animals were prepared as described above, but were incubated and recorded in ACSF containing 1 µM reserpine.

## Fiber photometry

Upon stereotaxic injection of AAV9-hSyn-GRAB_DA_2m_ in dorso-lateral striatum, a 400 μm optic fiber (FP400URT, Thorlabs) housed in a 1.25 mm ceramic ferrule (CFLC440, Thorlabs) was implanted 0.2 mm above the injection site using C&B metabond (Parkel). Following 2 weeks of recovery in their home cage, mice were habituated for a minimum of 5 days to locomote while head-fixed on a cylindrical wheel placed into a dark soundproof chamber and were water restricted to incite consumption of water rewards delivered from a spout at semi-random 8 to 12 s intervals. Licking was monitored by a capacitive touch sensor (AT42QT1010, Sparkfun). Extracellular DA levels in striatum were imaged by providing 470 nm excitation light from a LED (M470F3; Thorlabs) coupled to an optical fiber (400 μm, 0.48 NA; Doric) to a fluorescence mini-cube (FMC3_E(460-490)_F(500-550)_S; Doric) connected to the mouse via a fiber optic patch cord (400 μm, 0.48 NA; Doric) and zirconia sleeve. Emission light (500-550 nm) was collected through the same patch cord and mini-cube by a femtowatt photoreceiver (2151; Newport) fitted with a FC adapter (FOA_2151_FC; Doric) via a 600 μm 0.48NA fiber optic (Doric).

## Immunohistochemistry

Mice were deeply anesthetized with isoflurane and perfused transcardially with 4% paraformaldehyde in 0.1 M sodium phosphate buffer. Brains were post-fixed for 1–3 days, sectioned coronally (50 µm in thickness) using a vibratome (Leica; VT1000S) and processed for immunofluorescence staining for tyrosine hydroxylase (Sigma-Aldrich; AB152, 1:2000), Gat1 (Cell Signaling; 37342, 1:200), Aldh1a1 (R&D Systems; AF5869-SP, 1:200), HA (Abcam; ab18181, 1:500). After primary antibodies incubation (4°C, overnight), the following secondary antibodies (all Thermo Fisher Scientific) were applied for 2 hrs at room temperature: Goat anti-mouse IgG Alexa Fluor 568 (A11004), Goat anti-mouse IgG Alexa Fluor 488 (A11029), Goat anti-rabbit IgG Alexa Fluor 488 (A11034), Goat anti-rabbit IgG Alexa Fluor 568 (A11036), Donkey anti-goat IgG Alexa Fluor 568 (A11057), Donkey anti-rabbit IgG Alexa Fluor 488 (A21206). Brain sections were mounted on Superfrost slides and coverslipped with DAPI Fluoromount-G (SouthernBiotech, 0100-20).

## In situ hybridization

Mice were deeply anesthetized with isoflurane, the brain was rapidly extracted and frozen on dry ice in plastic cryomolds containing OCT compound (Fisher Scientific, 4585). Coronal sections (16 µm in thickness) were obtained using a Leica CM3050S cryostat, immediately mounted onto Superfrost Plus glass slides (Fisher Scientific, 1255015) and kept at -80°C until use. RNAscope assay was performed following Advanced Cell Diagnostic (ACD) manual Fluorescent Multiplex Kit. Briefly, slides were fixed in ice-cold 4% PFA for 15 min, then dehydrated in 50%, 70% and 100% ethanol before incubation with protease IV at room temperature for 30 min. The following target probes were used: *Slc6a3*/Dat (31544-C2), *Th*/TH (317621), *Gad1*/Gad67 (400951-C3), *Gad2*/Gad65 (439371-C2). For *Slc6a1*/Gat1, a custom probe was designed by ACD targeting 243-718 bp of NM_178703.4. After the last wash step, sections were incubated in DAPI at room temperature for 30 sec before coverslipping with Fluoromount-G (SouthernBiotech, 0100-01).

## Data acquisition and analysis

Membrane currents were amplified and low-pass filtered at 2 kHz using a Multiclamp 700B amplifier (Molecular Devices), digitized at 10 kHz using National Instruments acquisition boards (BNC 2110) and a custom version of ScanImage (Pologruto et al., 2003 https://github.com/bernardosabatinilab/SabalabSoftware_Nov2009) written in MATLAB (Mathworks). Electrophysiology data were analyzed offline using Igor Pro 6.02A (Wavemetrics). In figures, voltage-clamp traces represent the averaged waveform of 3–5 consecutive acquisitions. Single waveforms aligned to optogenetic light stimulus onset were used to measure postsynaptic current peak amplitude, latency (from light onset to current onset), and 10–90% rise time. Only oIPSCs and oEPSCs larger than 11 pA in amplitude were considered to compute prevalence, latency and rise time. For sIPSCs, the detection threshold was set to 20 pA to facilitate event detection in the presence of large and noisy tonic GABA currents. For photometry, signals were continuously digitized at 2 kHz using a National Instruments acquisition board (USB-6343) and Wavesurfer software (Janelia), low-pass filtered and down-sampled to 30 Hz, and aligned to the first lick within 500 ms of water delivery for analysis in MATLAB. For *in situ* hybridization and immunofluorescence analyses, brain sections were imaged with an Olympus VS120 slide-scanning microscope. High-resolution images of regions of interest were subsequently acquired with a Zeiss LSM 800 confocal microscope. Confocal images in figures represent maximum intensity projections of 3-μm confocal stacks. Cell bodies were segmentated with Cellpose (Stringer et al., 2021) using the TH signal as a mask and colocalization was quantified using ImageJ (NIH). Data (reported in text and figures as mean ± s.e.m.) were compared using Prism 9 (GraphPad) with the following non-parametric statistical tests (as indicated in the text): Mann–Whitney for comparisons between test and control groups, and Kruskal–Wallis analysis of variance (ANOVA) followed by Dunn’s Multiple Comparison Test for multiple group comparisons. Two-way repeated measures ANOVA were used in experiment characterizing the time course of synaptic transmission rundown. For electrophysiology experiments, n-values represent the number of recorded cells from N mice. In most cases, a single cell was recorded per slice, with each animal contributing at most 5 individual recordings per condition. For immunohistochemistry and in situ hybridization, n represents either the number of sections analyzed (with 3 sections analyzed per mouse, on average) or the total number of DA cells quantified across multiple sections in 3 or more mice per condition. For fiber photometry recordings, n represents the number of imaging sessions performed, with a maximum of 2 imaging sessions per mouse. Exact p-values are provided in text and figure legends, and statistical significance in figures is presented as * p < 0.05, ** p < 0.01 and *** p < 0.001.

---

**Figure S1.**
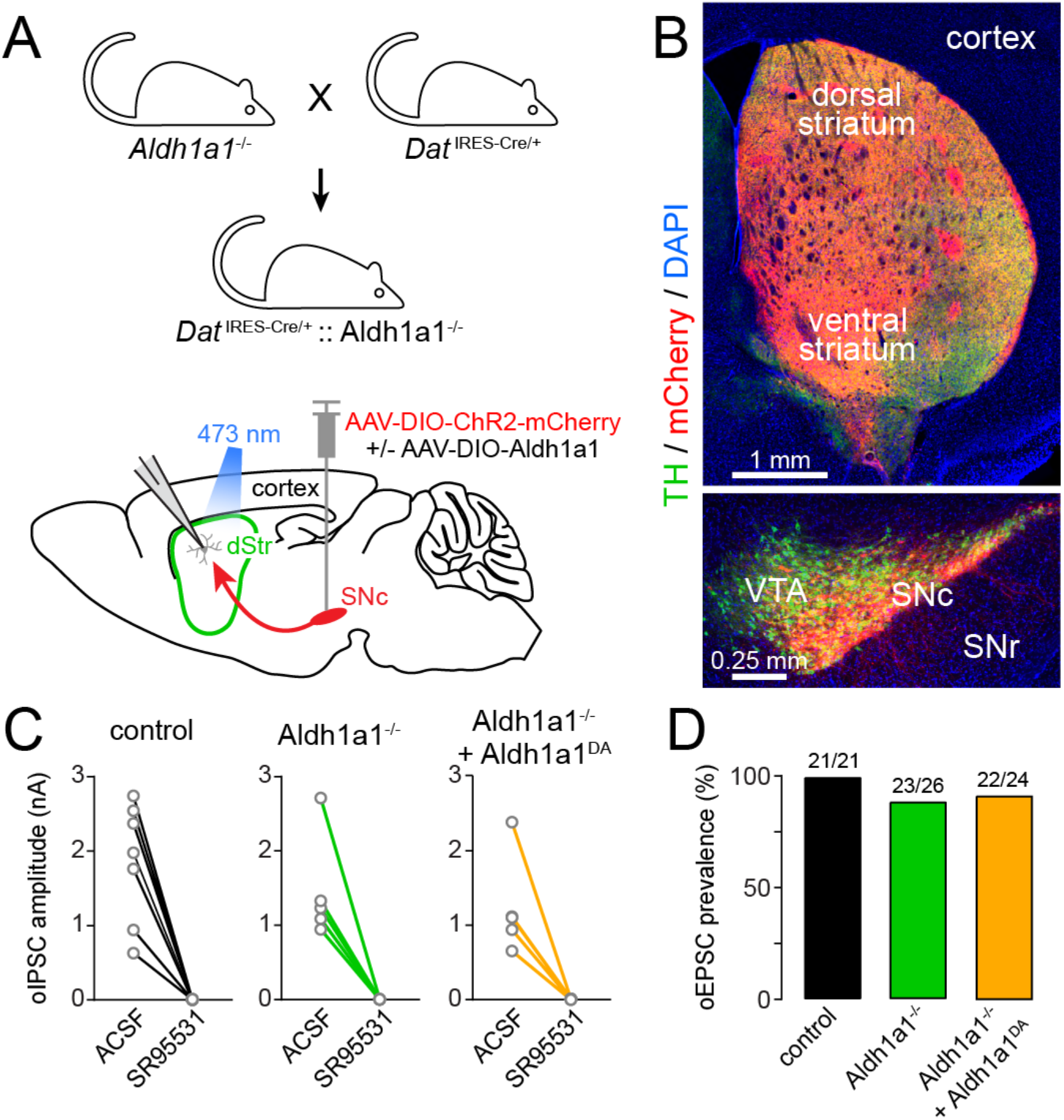
Experimental setup. **A)** Diagram of genetic strategy to generate mice in which the gene encoding Aldh1a1 is intact or knocked out, while gaining genetic access to midbrain DA neurons through intracranial injection of AAVs encoding Cre-dependent (DIO) ChR2-mCherry or Cre-dependent Aldh1a1. **B)** Coronal forebrain (top) and ventral midbrain (bottom) sections from a control *Dat*^IRES-Cre/+^ mouse expressing Cre-dependent ChR2-mCherry in midbrain DA neurons for 3 weeks immunolabeled for tyrosine hydroxylase (TH, green) and nuclear stained with DAPI (blue). ChR2-mCherry expression is limited to TH+ DA neurons in SNc/VTA and distributes throughout axons in striatum. **C)** Amplitude of optogenetically evoked IPSCs in SPNs before (ACSF) and after bath application of the GABAA receptor antagonist SR95531 (10 μM) in 3 different genotypes. **D)** Prevalence of oEPSCs exceeding our detection threshold of 11 pA in SPNs from 3 different genotypes. Genetically deleting Aldh1a1 does not affect our ability to stimulate DA axons or record postsynaptic responses.

**Figure S2.**
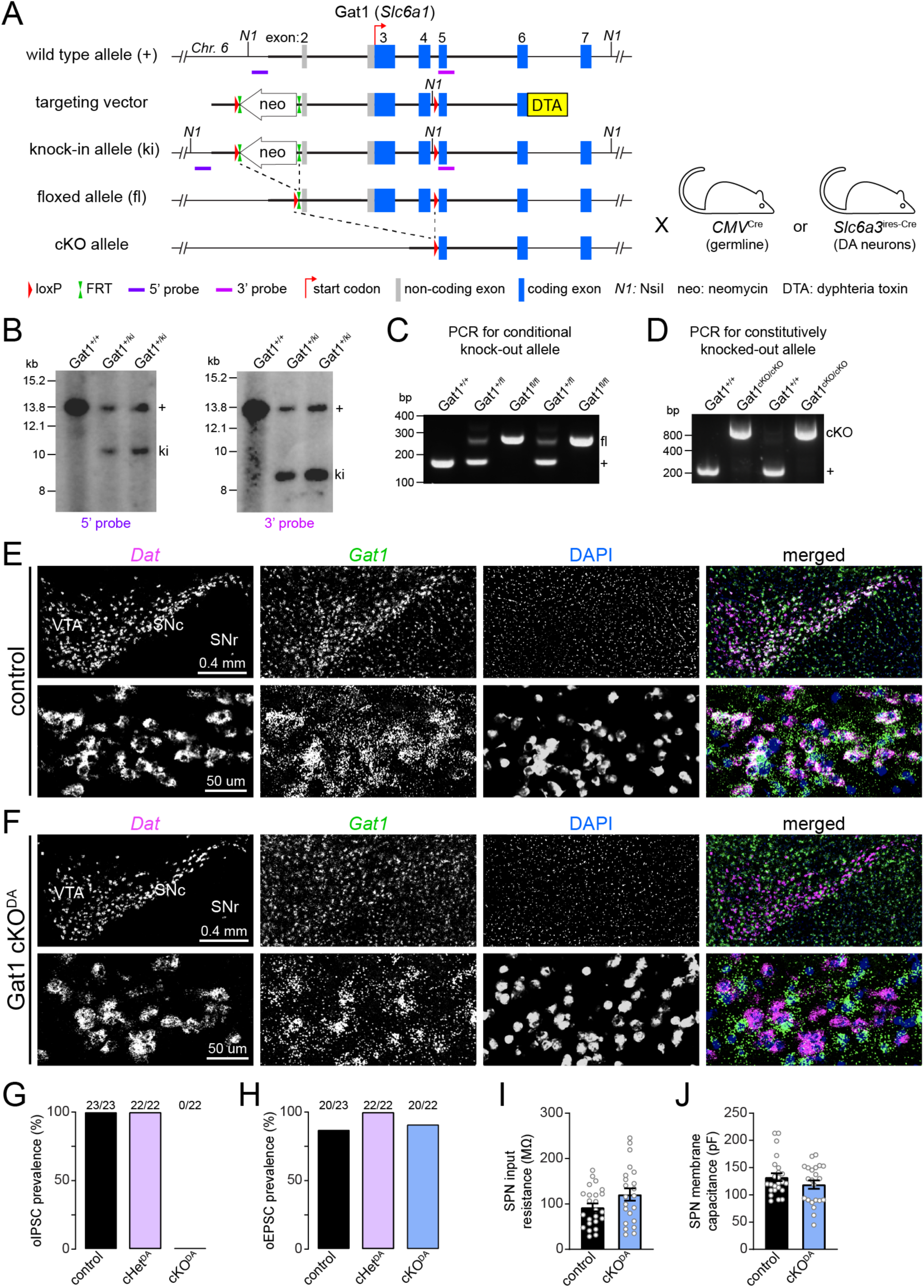
Generation and validation of Gat1 conditional knockout mice. **A)** Schematic of targeting strategy (not drawn to scale). Solid horizontal line represents relevant portion of *Slc6a1* genomic locus on mouse chromosome 6. Rectangles represent Slc6a1 exons (numbered), with coding sequences highlighted in blue and non-coding exons in gray (the initiation (ATG) codon is highlighted with red angled arrow). Thick horizontal lines depict intronic sequences included in targeting vector for homologous recombination. Targeting vector contains a neomycin positive selection cassette flanked by FRT sites, a diphteria toxin negative selection marker on 3’ end, two LoxP sites (one between exons 1 and 2, and another between exons 4 and 5) and a novel NsiI restriction site. ES cell clones with correct 5’ and 3’ recombination events (i.e. knockin allele) were injected into blastocyst-stage embryos and implanted in pseudo-pregnant females to generate chimeric males, which were bred to Flp-deleter mice to excise the Neomycin cassette and generate mice heterozygous for the conditional knockout allele. Cre recombinase subsequently excises exons 2-4 to functionally delete Slc6a1 only in Cre-expressing cells. **B)** Southern blot analysis for correct 5’ (left) and 3’ (right) homologous recombination in ES cells. NsiI-digested genomic DNA from two ES cell clones with recombined ki alleles show expected restriction fragments for one wild-type allele (13.7 kb) and one ki allele (10.2 kb for 5’ hybridization probe and 8.3 kb for 3’ hybridization probe). Wild-type DNA is shown in first lane for comparison. **C)** Example PCR gel for Gat1 conditional knockout mice (prior to Cre-mediated recombination) with GAT1-FloxF and GAT1-FloxR primers yielding expected wild-type (170 bp) and floxed-allele (276 bp) bands in 5 littermates. **D)** Same as (**C**) for Gat1^fl^ mice after germline Cre-mediated recombination (using *CMV*^Cre^ mice) and *Gat1*^fl^ mice, which effectively yields a conventional constitutive Gat1 knockout allele, using GAT1-Flox14F, GAT1-Flox18F and GAT1-Flox15R primers to obtain expected wild-type (211 bp) wild-type and recombined/knockout (818 bp) bands. **E)** Representative low (top) and high magnification (bottom) epifluorescence images of fluorescence *in situ* hybridization for *Dat* (magenta) and *Gat1* (green) in coronal section of ventral midbrain of a control mouse (*Dat*^IRES-Cre/+^;*Gat1*^+/+^). SNc is visible with *Gat1* antisense probe alone. Colocalization with DA cell bodies appears white in merged images. **F)** Same as (**E**) for Gat1 cKO^DA^ mouse (*Dat*^IRES-Cre/+^;*Gat1*^fl/fl^). Note how *Gat1* signal remains intact everywhere except for DA neurons; SNc is no longer visible with *Gat1* antisense probe alone and colocalization is no longer evident in merged images. **G–H)** Percentage of all recordings showing a ChR2-evoked IPSC (**G**) or EPSC (**H**) from SNc^DA^ axons in dorsal striatum SPNs in control, Gat1 cHet^DA^ and Gat1 cKO^DA^ slices. Number of recordings per condition is indicated above each bar. **I)** Input resistance of SPNs recorded in control and Gat1 cKO^DA^ slices (p = 0.15, Mann–Whitney ). Data show Individual values along with mean ± s.e.m. **J)** Same as (**I**) for membrane capacitance (p = 0.42, Mann–Whitney ).

**Figure S3.**
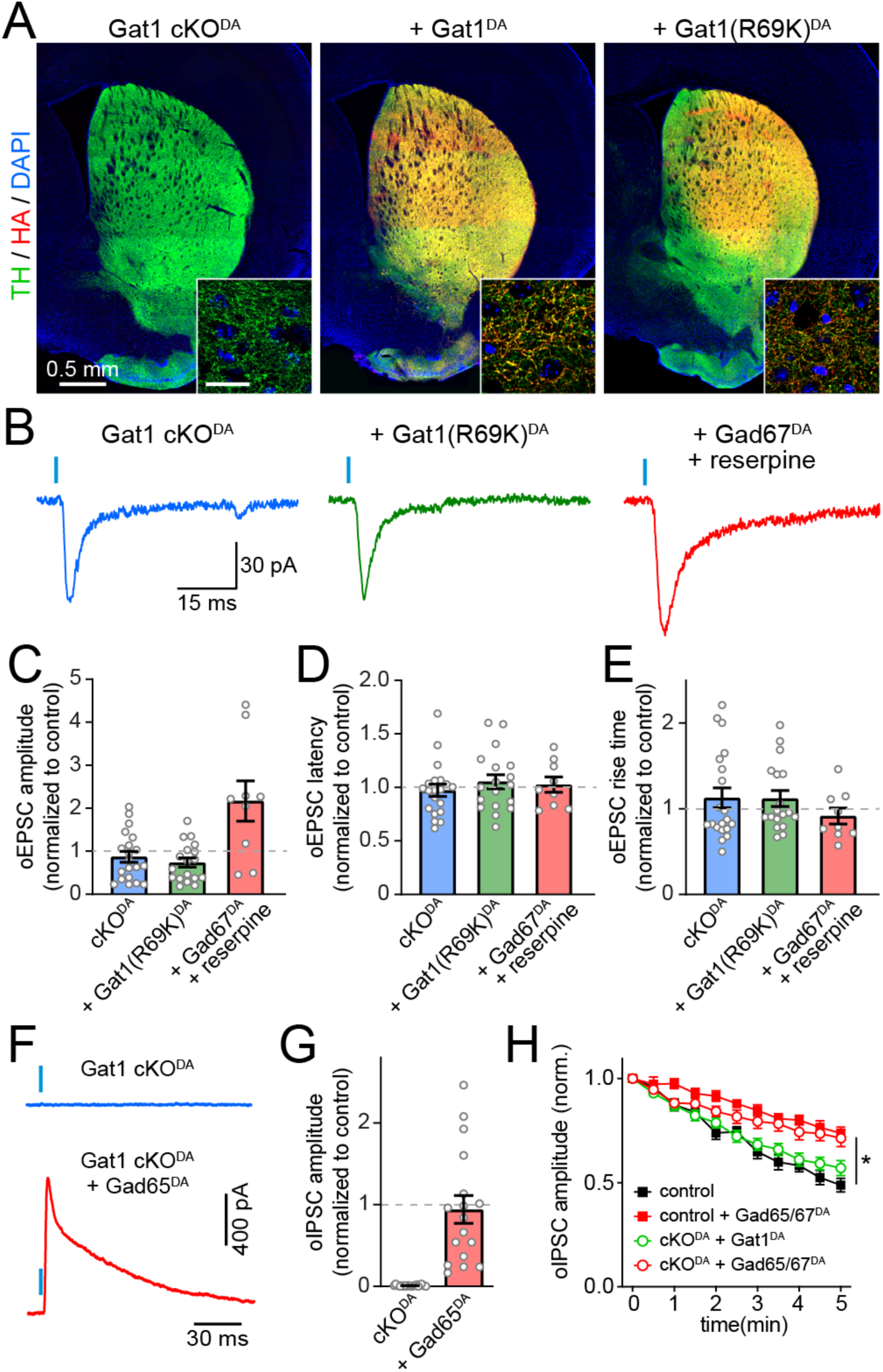
Controls for ChR2-mediated activation of SNc^DA^ axons across conditions. **A)** Epifluorescence images of coronal forebrain sections immunostained for TH (green) to label DA axons, HA (red) to label constructs exogenously-expressed in DA neurons, and DAPI (blue) to label nuclei in Gat1 cKO^DA^ mice (*Dat*^IRES-Cre/+^;*Gat1*^fl/fl^) injected in ventral midbrain with AAVs encoding Cre-dependent ChR2-mCherry alone (left), ChR2-mCherry + Gat1-HA (middle), and ChR2-mCherry + Gat1(K69K)-HA (right). Insets: high magnification view in dorsal striatum (scale bar: 20 μm). Note Gat1’s distribution in axons. **B)** Example EPSCs evoked by optogenetic stimulation of SNc^DA^ axons in dorsal striatum SPNs in slices from Gat1 cKO^DA^ mice (blue) expressing Gat1(R69K) (green) or Gad67 (red) in DA axons. **C–E)** Quantification of oEPSC amplitude (**C**), latency (**D**) and rise time (**E**) normalized to *Dat*^IRES-Cre/+^ controls. **F)** Representative IPSCs recorded from Gat1 cKO^DA^ SPNs held at 0 mV upon optogenetic stimulation of DA axons overexpressing Gad65 (red) or not (blue). **G)** oIPSC amplitude normalized to Dat^IRES-Cre/+^ controls (Gat1 cKO^DA^+Gad65^DA^: n = 17 SPNs from 5 mice; p = 0.99 vs. control, p = 3.6 x 10^-7^ vs. Gat1 cKO^DA^; Dunn’s Multiple Comparison). **H)** Quantification of oIPSC amplitude vs. time in control slices (filled squares) and Gat1 cKO^DA^ (empty circles) expressing either Gat1 (green) or Gad65/Gad67 (red) in SNc^DA^ neurons. Gat1 cKO^DA^ + Gat1^DA^ (n = 14 SPNs in 4 mice), Gat1 cKO^DA^ + Gad65/67^DA^ (n = 25 SPNs in 8 mice). In control mice, the amplitude of oIPSCs decreased significantly less when SNc^DA^ neurons expressed Gad65 or Gad67 (control: n = 22 SPNs from 9 mice; Gad65/67: n = 17 SPNs from 6 mice; two-way ANOVA group x time: F_10,370_ = 10.55, p = <0.0001). In Gat1 cKO^DA^ mice too, oIPSCs were less prone to rundown when SNc^DA^ express Gad65 or Gad67 (n = 25 SPNs from 8 mice) vs. Gat1 (n = 14 SPNs from 4 mice; two-way ANOVA group x time: F_10,365_ = 3.15, p = 7 x 10^-4^).

## Notes

### Competing Interest Statement

The authors have declared no competing interest.

